# OPTN protects retinal ganglion cells and ameliorates neuroinflammation in optic neuropathies

**DOI:** 10.1101/2025.01.09.632264

**Authors:** Qinglong Wang, Yiqi Wang, Yi Da Douglas Jiang, Ryan Donahue, Gaby Cao, Weixuan Yan, Hong Guo, Jin Hao, Yi Lu, Qianbin Wang, Feng Tian

**Affiliations:** Department of Neurology, Beth Israel Deaconess Medical Center, Harvard Medical School, Boston, MA 02115, USA; Department of Biology, Georgia Institute of Technology, Atlanta, GA 30332, USA; F.M. Kirby Neurobiology Center, Boston Children’s Hospital, and Department of Neurology, Harvard Medical School, 300 Longwood Avenue, Boston, MA 02115, USA; School of Arts and Sciences, Tufts University, Medford, MA 02155, USA; Department of Neurosurgery, Duke University Medical Center, Durham, NC 27710, USA; Department of Neurosurgery, Brigham and Women’s Hospital, Boston, MA 02115, USA; Department of Stem Cell and Regenerative Biology, Harvard University, Cambridge, MA 02138, USA; Department of Biomedical Engineering, Binghamton University, State University of New York, Binghamton, NY 13902, USA

## Abstract

Optineurin (OPTN) is a crucial component of the homeostatic pathway, playing a pivotal role in regulating a number of essential signaling pathways including NF-κB, interferon, autophagy, and vesicular trafficking. The dysfunction of OPTN has been implicated in the pathogenesis of several diseases, such as primary open angle glaucoma (POAG), amyotrophic lateral sclerosis (ALS), frontotemporal lobar dementia, and Paget’s disease of bone. Interestingly, mutations in OPTN are implicated as gain-of-function in glaucoma pathology and loss-of-function in ALS. However, the role of loss-of-function OPTN in glaucoma pathology remains unclear. Here, we demonstrate that OPTN dysfunction contributes to chronic neuroinflammation, leading to sustained RGC death, which may represent a shared pathological mechanism in both normal tension glaucoma (NTG) and high-tension glaucoma (HTG). Retinal conditioned OPTN knockout contributes to short-term astrogliosis and long-term microglia activation, with the propagation of microglia activation spreading to the optic nerve. Moreover, OPTN loss of function does not further exacerbate RGC death in the ocular hypertension mouse model induced by viscobead injection. Combined with the downregulation of OPTN in glaucoma patients, we have revealed an IOP-independent mechanism of glaucoma pathogenesis. Furthermore, we found that OPTN-driven NPY upregulation may suppress the CHOP-associated neurodegeneration. Our findings reveal a neuroprotective role for the OPTN-NPY signaling pathway, and its dysfunction promotes RGC loss in glaucoma pathology. The OPTN-NPY-mediated neuroinflammatory pathway provides a potential therapy for IOP-resistant glaucoma and highlights a druggable target for CHOP-associated neurodegeneration.

---

Optineurin (OPTN) is an important adaptor protein involved in several critical signaling pathways including neuroimmune homeostasis^1^, protein trafficking, and organelle maintenance^2^. Mutations of OPTN have been implicated in the pathogenesis of several diseases, such as primary open-angle glaucoma (POAG), amyotrophic lateral sclerosis (ALS), frontotemporal lobar dementia, and Paget’s disease of bone^3^. Retinal ganglion cells (RGCs) are retinal neurons that transmit visual input from the eye to the brain. Degeneration of RGCs and their axons in the optic nerve are cardinal features of glaucoma and other optic neuropathy conditions, which lead to irreversible vision loss^4–7^. Importantly, among the genetic mutations associated with glaucoma, mutations in OPTN, were found in 16.7% of families with hereditary POAG, including individuals with normal intraocular pressure (IOP) or normal tension glaucoma (NTG)^8^. The wild-type OPTN plays a crucial role in multiple cellular processes, including autophagy, endocytic trafficking, NF-κB regulation, neuroinflammation, and transcriptional activation^9–13^. Contrary to the disease-related OPTN mutations in ALS, which act in a loss-of-function manner^14^, published studies on glaucoma-associated OPTN mutations seem to suggest a gain-of-function pattern in optic neuropathies^15,16^. Intriguingly, *In vitro* studies have shown that suppression of OPTN expression in cultured primary RGCs and RGC-5 cells suppresses cell growth, increases apoptosis, and reduces neurotrophic support, suggesting a potential neuroprotective role of wild-type OPTN^12,17^. However, direct evidence related to the role of wild-type OPTN in RGC homeostasis and degeneration *in vivo* remains unclear. In addition, despite evidence supporting a potential neuroprotective function during RGC development^18^, it is unclear whether OPTN also causes POAG via a loss-of-function mechanism. Therefore, a comprehensive understanding of the neurodegenerative signaling pathways associated with OPTN dysfunction would greatly facilitate the dissection of the shared and distinct mechanisms among multiple neurodegenerative conditions, especially NTG, high-tension glaucoma (HTG), and ALS.

While the patterns of optic nerve damage and visual field impairment somewhat differ between NTG and HTG^19,20^, they share substantial pathological similarities, including activation of NF-κB signaling, elevated oxidative endoplasmic reticulum (ER) stress, RGC apoptosis, and gliosis^21–27^, all of which are closely associated with the physiological function of wild-type OPTN^9–13,28^. Meanwhile, it has been suggested that in NTG, changes in RGC susceptibility may lead to their death within the normal range of IOP^29,30^. Lowering IOP is also beneficial for NTG to some extent^31^. Notably, even with significant IOP reduction, approximately 50% of POAG patients continue to experience progressive visual field loss^32^, highlighting the importance of IOP-independent therapeutic strategies. As one of the most prevalent disease-associated genes in NTG, how OPTN loss of function contributes to RGC susceptibility and pathogenesis of NTG and HTG is a critical but unanswered question.

In addition to the intrinsic susceptibility of RGCs to glaucomatous pathology, the responses of glial cells play a significant role in determining neuronal outcomes. Both astrocytes and microglia respond robustly to the neurodegenerative stress in optic neuropathies^33^. Chronic neuroinflammation, as a result of the activation of these glial cells, is increasingly recognized as a key factor in optic neuropathies and various neurodegenerative conditions^34–36^. In a microbead occlusion model of glaucoma, astrocytes have been demonstrated to transform into a neurotoxic reactive state^37,38^. Microglia, functioning as the primary innate immune cells in the central nervous system (CNS), play a pivotal role in the neuroimmune homeostasis^39^. Upon retinal or optic nerve damage, they may transition to an activated state and migrate to sites of injury, potentially contributing to RGC death by releasing neurotoxic factors^40,41^. Microglia activation is hypothesized in the field as a significant contributing factor to retinal neurodegeneration, which might explain why IOP reduction cannot fully prevent progressive RGC death in glaucoma^42^. Resetting activated microglia to a homeostatic state through IGFBPL1 administration can alleviate neuroinflammation and prevent neurodegeneration and visual loss in glaucoma^36^. In addition, key transcription factors (TFs) involved in RGC degeneration, such as CHOP and ATF3^43–45^, promote neurodegeneration through the activation of pro-inflammatory signaling pathways^43,46–49^. Therefore, targeting neuroinflammation and preserving neuroimmune homeostasis represent promising neuroprotective strategies for the treatment of glaucoma and potentially other neurodegenerative conditions^34^. However, despite considerable attention given to the role of inflammation and innate immune response^50,51^, it remains unclear whether OPTN plays a role in neuroimmune homeostasis and RGC vulnerability in neurodegeneration.

Here, we asked the question of whether wild-type OPTN is essential for regulating RGC homeostasis and vulnerability in neurodegeneration, and whether OPTN dysfunction represents a shared pathological mechanism underlying both NTG and HTG. Our findings demonstrate that wild-type OPTN plays a pivotal role in maintaining RGC survival and regulating neuroimmune homeostasis in the retina. This IOP-independent pathway mediated by OPTN may be shared between NTG and HTG. These results underscore the significance of restoring OPTN-mediated neuroimmune homeostasis to impede retinal neurodegeneration.

## Results

### OPTN loss of function leads to chronic RGC loss without IOP elevation

To examine the effects of OPTN deficiency on retinal homeostasis, we performed the intravitreal injection of AAV2-Cre and AAV2-PLAP (control group) to 1-month-old OPTN floxed mice (*Optn^tm1.1Jda^*/J; Jackson Labs; Stock# 029708), resulting in a loss-of-function mutation of OPTN with C-terminus truncation (starting from exon 12) (Fig. 1a)^52,53^. Using immunostaining of retinal sections for the RGC marker RBPMS and OPTN C-terminus truncation protein, we detected that the percentage of OPTN expressing RGCs was significantly decreased at 1 month post intravitreal injection of AAV2-Cre, but not in AAV2-PLAP injected retinas (Supplementary Fig. 1a-b). IOP measurements were performed weekly after 4 weeks post injection. Retinas and optic nerves were collected at 1-, 3-, and 6-month post injection for immunohistochemical (IHC) analysis. To measure the RGC loss after OPTN loss-of-function, we counted the RBPMS+ RGCs in AAV2-Cre and AAV2-PLAP groups compared with those in the intact retina through sectioned and whole mount retinas (Fig. 1b, Supplementary Fig. 1c). We demonstrated that RGC survival progressively decreased at 1-, 3-, and 6-month post AAV2-Cre injection, whereas it remained unchanged in AAV2-PLAP injected controls (Fig. 1c). Therefore, OPTN plays a role in maintaining RGC survival, and its deficiency leads to the progressive death of RGCs. Moreover, the knockout of OPTN did not cause any changes in IOP, which is consistent with its role in NTG (Fig. 1e).

**Fig. 1.**
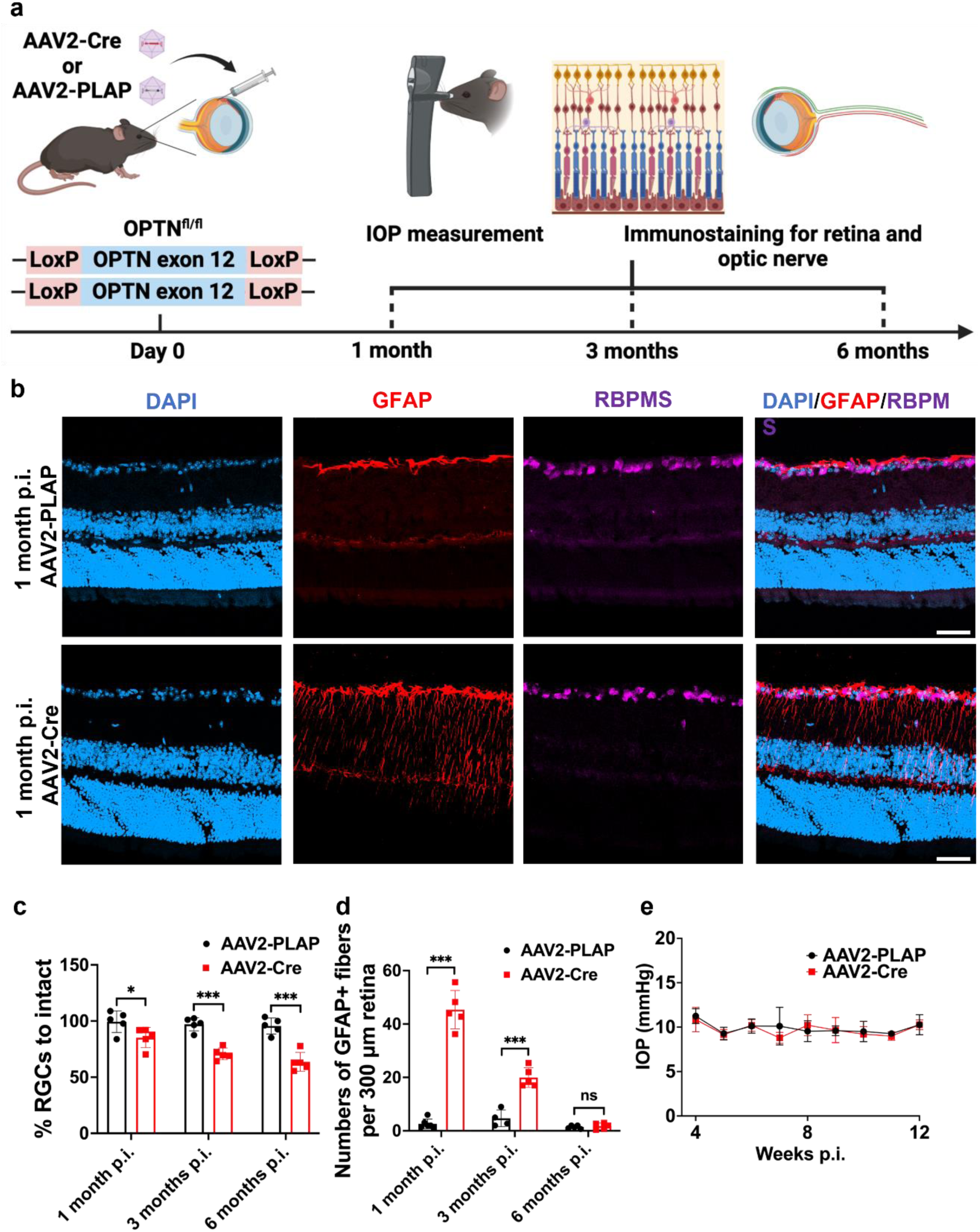
OPTN loss-of-function leads to RGC loss and astrogliosis. **a** Schematic illustration of the OPTN ablation in retina. **b** Representative IHC images showing OPTN knockout induced RGC loss and astrogliosis 1, 3, and 6 months post intravitreal injection. Scale bar, 50 μm. **c,d** Quantification of RGC survival (**c**) and astrogliosis (**d**) with OPTN knockout 1, 3, and 6 months post intravitreal injection. Data are shown as mean ±s.e.m. with n = 5 biological replicates. \**p* < 0.05, \*\**p* < 0.01, \*\*\**p* < 0.001, ns, not significant, calculated by unpaired t-test. **e** Intraocular pressure (IOP) of mice post intravitreal injection of AAV2-Cre or PLAP virus shows no significant difference between OPTN knockout and control mice. Data are shown as mean ±s.e.m. with n = 10.

### OPTN loss-of-function results in short-term reactive astrogliosis in the retina

As important cells for maintaining tissue homeostasis in the retina and optic nerve, astrocytes are reactivated under glaucoma pathology. In glaucomatous astrocytes, there is an upregulation of glial fibrillary acidic protein (GFAP), an enlargement of the cytoskeleton, and elongated cellular processes^33^. Along with RGC loss, we observed significant reactive astrogliosis, which is shown by the astrocytic fiber count^54^ using immunolabeled with astrocyte marker GFAP, while astrogliosis was barely detectable in the control groups (Fig. 1b). We demonstrated that the reactive astrogliosis following AAV2-Cre injection was reversible, with a peak at 1-month post injection and returning to levels comparable to the control group by 6-month post injection (Fig. 1b, 1d, Supplementary Fig. 2). This suggests that OPTN dysfunction contributed to a short-term astrocytic response, with subsequent gradual resolution of astrogliosis over time. Given the ongoing death of RGCs at 6-month post injection, astrogliosis may not be the primary contributor to RGC loss after OPTN loss-of-function.

### OPTN loss-of-function leads to long-term microglia activation in the retina

Microglia activation is an early pathological hallmark of glaucoma-associated neurodegeneration and has been observed to release pro-inflammatory cytokines, leading to damage in RGCs across various animal models of glaucoma^42,55^. To further identify the role of OPTN in retinal homeostasis, we examined the microglia activation following OPTN knockout (Fig. 2a). Microglia are intrinsic regulators of homeostasis and key contributors to neuroinflammation^39^. CD68 was used to mark activated microglia in a phagocytic state^56,57^. We observed a basal level of microglia activation in the AAV2-PLAP injected retina, consistent with findings from a previous study^58^. However, following OPTN knockout, the proportion of activated microglia was significantly elevated compared to the control mice. This elevation persisted up to 6-month post injection (Fig. 2b). This suggests that OPTN knockout may promote the persistent and irreversible activation of microglia in the retina. Microglia morphology (Fig. 2c) was examined to accurately assess the impact of OPTN loss-of-function on their state transition. The results of microglia count analysis indicate that OPTN loss-of-function did not significantly alter microglia cell numbers (Fig. 2d). Morphological analysis of the microglia revealed the cell bodies enlarged (Fig. 2e), and their processes shortened (Fig. 2f), suggesting that the microglia exhibited a transformation into ameboid morphology following OPTN knockout. Together, OPTN deficiency leads to a persistent neuroinflammatory response that is difficult to resolve.

**Fig. 2.**
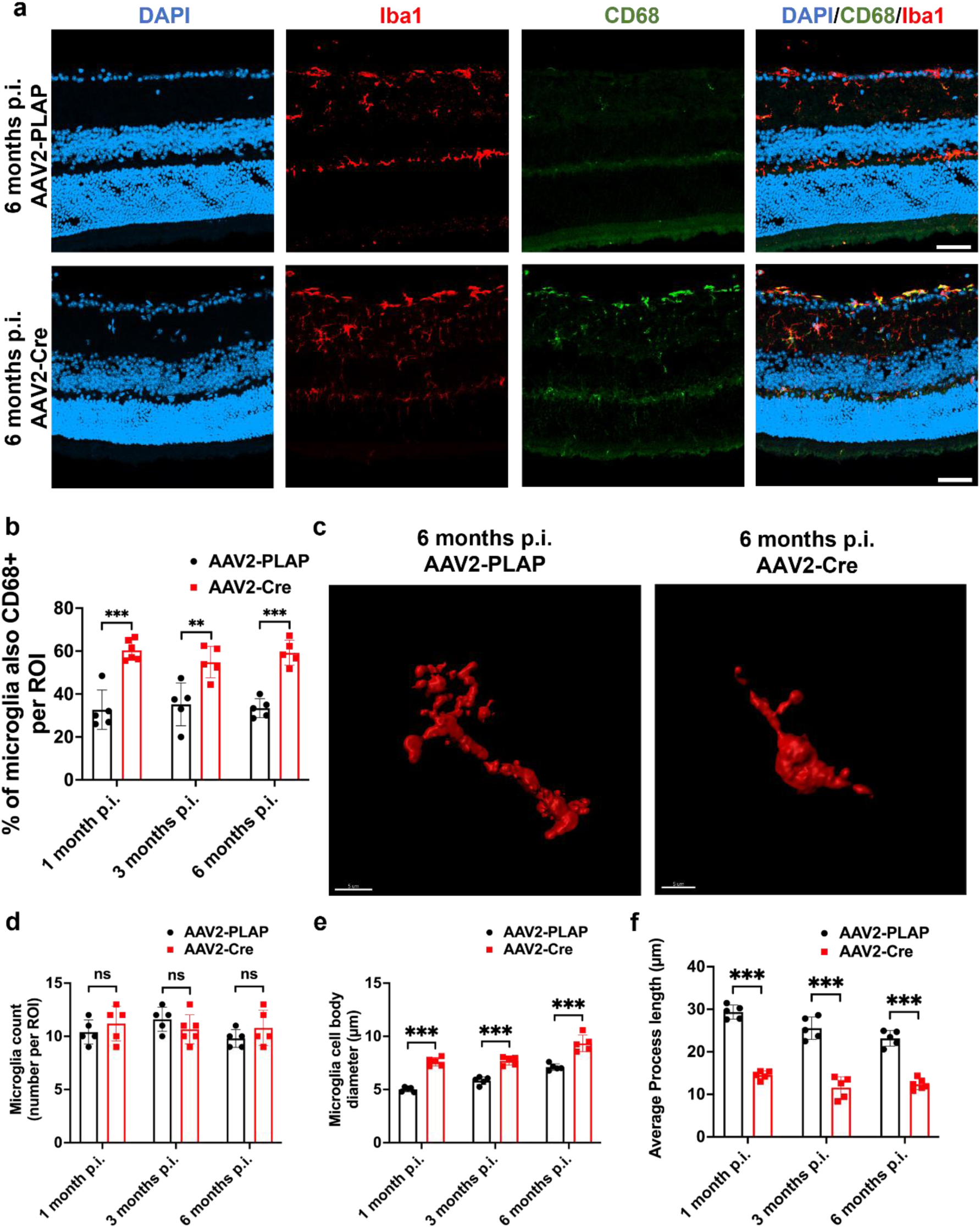
OPTN loss-of-function leads to microglia activation. **a** Representative IHC images showing OPTN knockout induced microglia activation 6 months post intravitreal injection. Scale bar, 50 μm. **b** Quantification of percentage of Iba1-positive microglia that are also CD68-positive with OPTN knockout 1, 3, and 6 months post intravitreal injection. Data are shown as mean ±s.e.m. with n = 5-6 biological replicates. \*\*\**p* < 0.001, ns, not significant, calculated by unpaired t-test. **c** A lateral view of 3D-reconstituted section retina with OPTN knockout 3 months post intravitreal injection. Scale bar, 5 µm. **d,e,f** Quantification of the number of Iba1-positive microglia (**d**), Iba1-positive microglia cell body diameter (**e**), and the average primary process length of Iba1-positive microglia (**f**) at 1, 3, and 6 months post intravitreal injection. Data are shown as mean ±s.e.m. with n = 5 biological replicates. \**p* < 0.05, \*\**p* < 0.01, \*\*\**p* < 0.001, ns, not significant.

### OPTN loss of function leads to microglia activation without astrogliosis in optic nerve

To investigate whether glaucomatous pathology caused by retinal OPTN loss-of-function, extends to the optic nerve, we conducted immunostaining to detect neuroinflammation at 3-month post OPTN knockout. Our findings indicate that OPTN knockout leads to activation of microglia (Fig. 3c, 3d) but no astrogliosis (Fig. 3a, 3b) in the optic nerve. This finding suggests that retinal OPTN may be involved in modulating the signaling pathways associated with microglia activation and that the neuroinflammatory response induced by OPTN knockout is spread to the optic nerve.

**Fig. 3.**
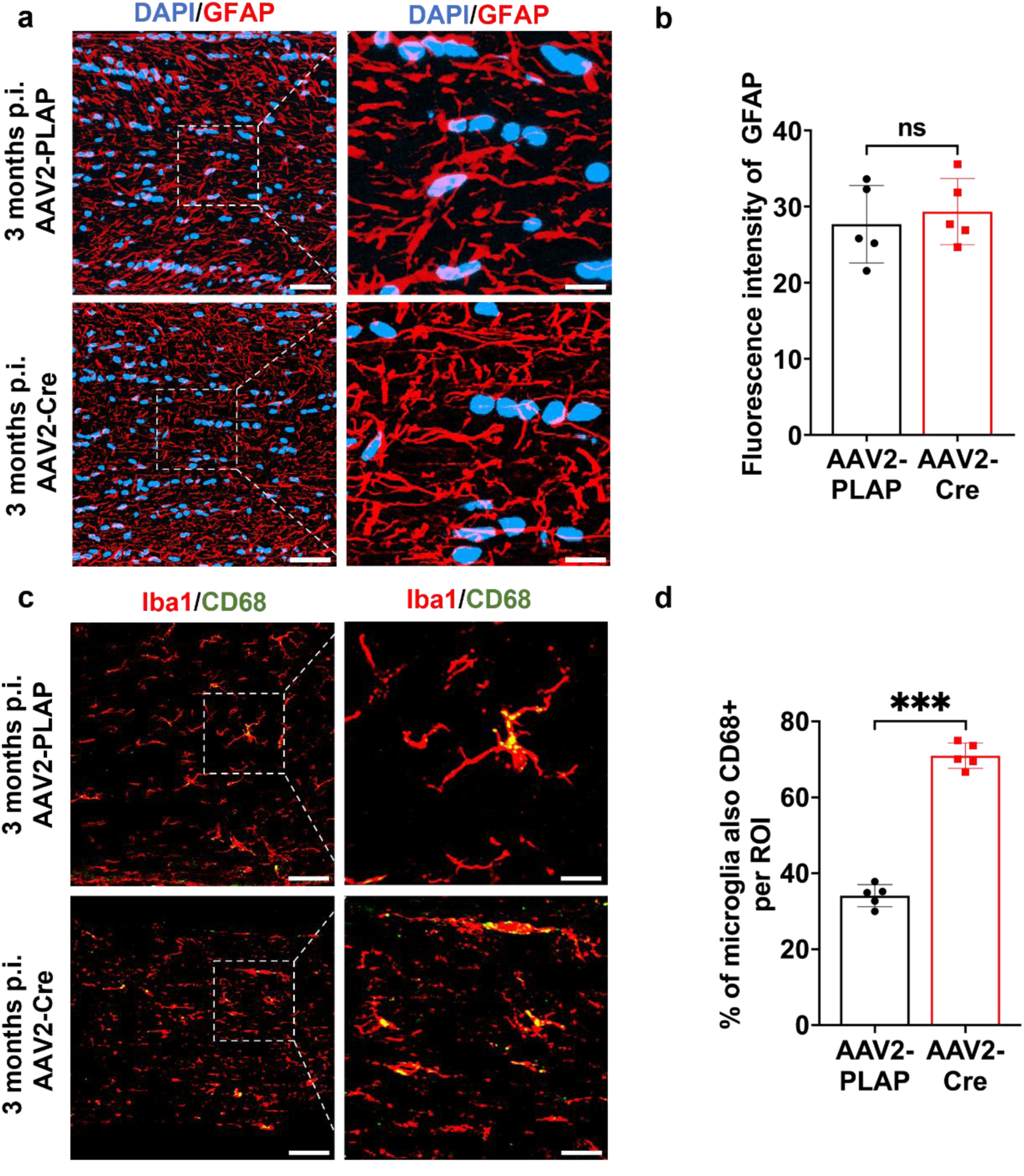
OPTN loss-of-function leads to microglia activation in optic nerve. **a, c** Representative IHC images showing OPTN knockout induced astrogliosis (**a**), and microglia activation (**c**) in optic nerve. Scale bar, 50 μm. **b, d** Quantification of astrogliosis (**b**), and microglia activation (**d**) in optic nerve 3 months post intravitreal injection. Data are shown as mean ±s.e.m. with n = 5 biological replicates. \*\**p* < 0.01, \*\*\**p* < 0.001, ns, not significant, calculated by unpaired t-test.

### OPTN loss-of-function does not further exacerbate RGC death in the ocular hypertension mouse model

The retinal OPTN knockout exhibited characteristic pathological features observed in NTG. We then introduced an HTG modeling method to analyze the role of the OPTN pathway in HTG pathology. Our recently developed viscobead injection model of hypertension glaucoma provides a more stable blockage of aqueous humor outflow with biodegradable viscoelastic beads and ocular hypertension^59^. We performed the viscobead injection 1 month after OPTN knockout, and the retinas were collected 3 months post-OPTN knockout (Fig. 4a). Concentrated viscobead were injected into the mouse anterior chamber to block aqueous drainage (Fig. 4c). During the 2 months following the viscobead injection, IOP remained significantly elevated, compared to the control group (Fig. 4d). IHC results showed that RGC survival rates following viscobead injection were similar in the OPTN intact group and the OPTN knockout group, with no significant difference (Fig. 4b, 4e). The control group (PBS injected) showed no significant difference in RGC survival compared to intact, demonstrating that the injection surgery will not result in the glaucomatous phenotypes. Ocular hypertension resulted in a greater loss of RGCs compared to the loss observed 3-month after OPTN loss-of-function. However, OPTN loss-of-function did not exacerbate RGC death under induced ocular hypertension, which implies that elevated IOP and OPTN loss of function may affect an overlapped group of vulnerable RGCs. This observation is consistent with clinical studies comparing HTG and NTG^60^. It could also be possible that some overlapped gene regulatory pathways might be present in regulating ocular hypertension and OPTN loss-of-function induced optic neuropathies.

**Fig. 4.**
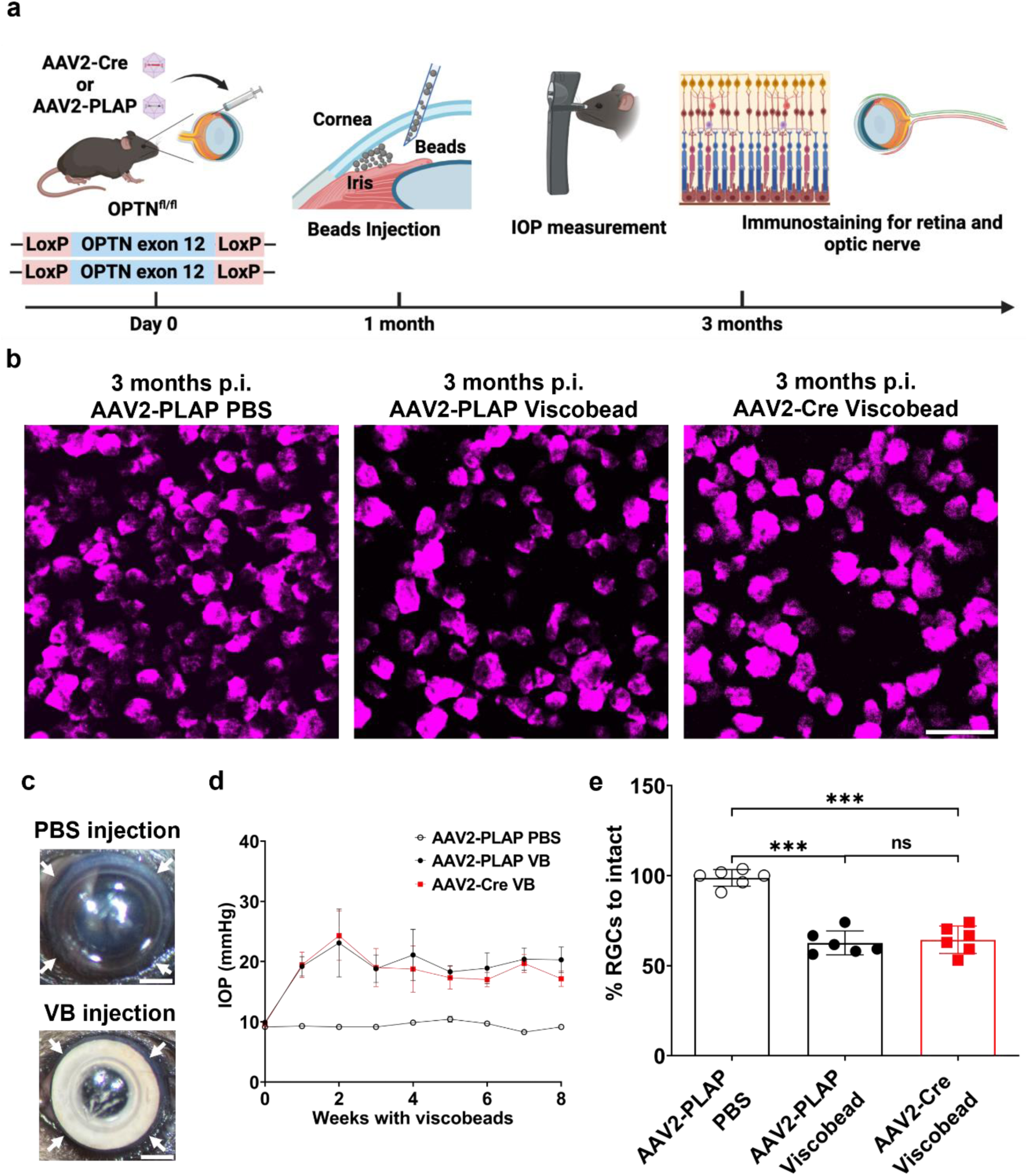
OPTN knockout and the high IOP model resulted in RGCs death in the same subgroup. **a** Schematic illustration of the viscobead injection model. **b** Representative IHC images showing viscobead injection induced RGCs loss in both OPTN knockout and control retina. Scale bar, 50 μm. **c** Representative photographs of viscobeads accumulated at mouse iridocorneal angle 5 min after injection. Arrows show that the white viscobeads were restricted at the iridocorneal angle. Scale bar, 1 mm. **d** IOP in the mice after the injection of viscobeads (n=10) or PBS (n=6) in OPTN knockout and control group. Data are shown as mean ±s.e.m.. **e** Quantification of RGC survival in different groups of the viscobead injection model. Data are shown as mean ±s.e.m. with n = 6 biological replicates. \**p* < 0.05, ns, not significant, calculated by unpaired t-test.

### OPTN loss-of-function and ocular hypertension induced the same extent of astrogliosis and microglia activation in the retina

OPTN loss-of-function resulted in RGC death and activation of neuroinflammatory pathways, independent of IOP. However, the involvement of OPTN-mediated neuroinflammatory pathways in the pathogenesis of glaucoma with high IOP remains unknown. To investigate whether the extent of neuroinflammation regulated by OPTN knockout is comparable to that induced by ocular hypertension, we quantified the astrogliosis and microglia activation provoked by high IOP in both OPTN intact and knockout retinas. The PBS-injected control group exhibited comparable levels of astrogliosis (Fig. 5a, 5b) and microglia activation (Fig.6a, 6b) to the OPTN intact control group (Fig.1d and Fig. 2b), indicating that the neuroinflammatory responses induced by surgical injury had resolved within 2 months. Along with RGC death, no significant difference in astrogliosis (Fig. 5a, 5b) and microglia activation (Fig. 6a, 6b) were observed in the OPTN knockout and intact groups following viscobead injection. These findings suggest that OPTN-mediated neuroinflammatory pathways may contribute to the neuroimmune response induced by ocular hypertension.

**Fig. 5.**
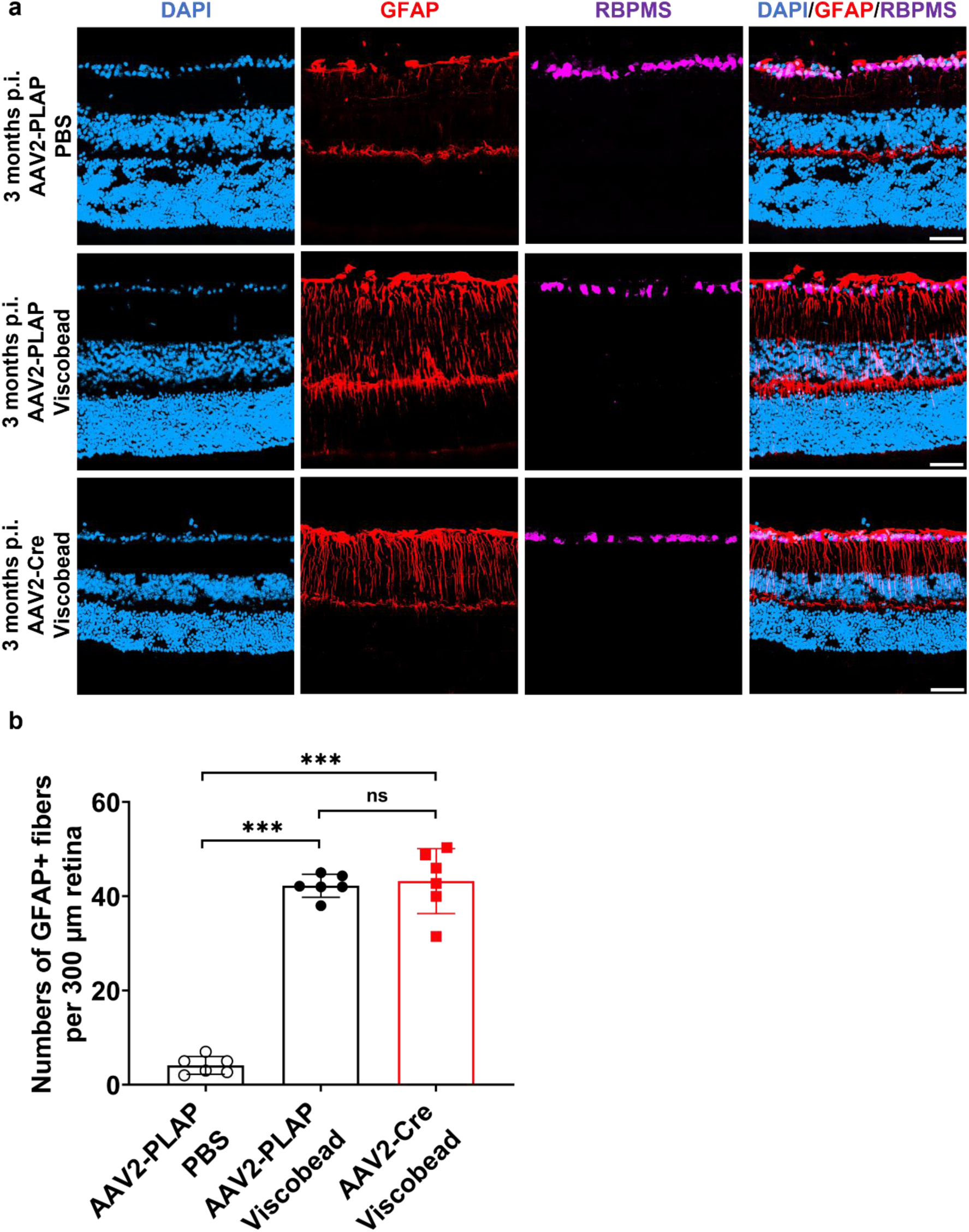
OPTN knockout and the high IOP model resulted in astrogliosis shown in retina. **a** Representative IHC images showing OPTN knockout induced astrogliosis. Scale bar, 50 μm. **b** Quantification of astrogliosis with OPTN knockout in retina. Data are shown as mean ±s.e.m. with n = 6 biological replicates. \*\*\**p* < 0.001, ns, not significant, calculated by unpaired t-test.

**Fig. 6.**
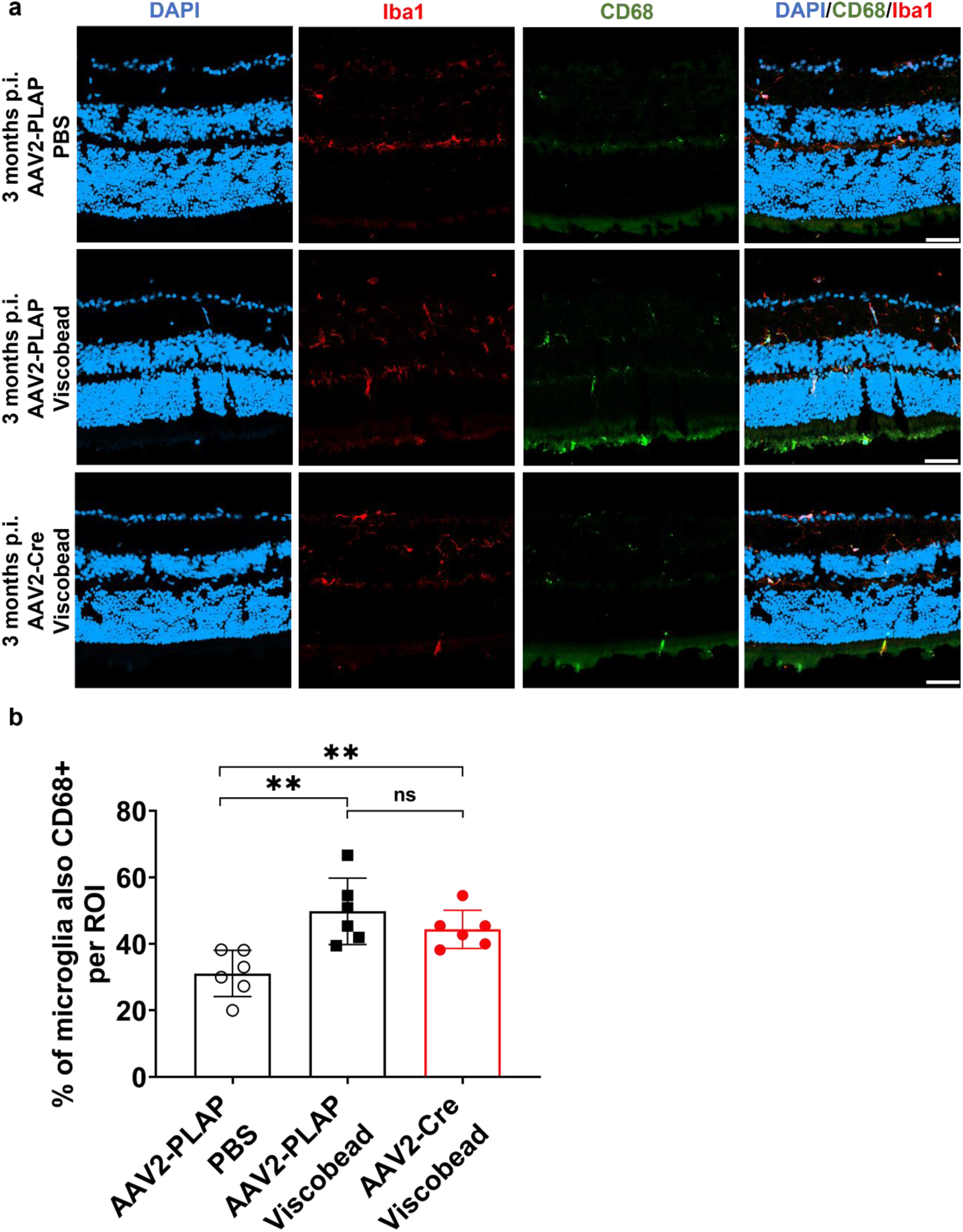
OPTN knockout and the high IOP model resulted in microglia activation shown in retina. **a** Representative IHC images showing OPTN knockout and the high IOP model induced microglia activation. Scale bar, 50 μm. **b** Quantification of microglia activation with OPTN knockout and the high IOP model in retina. Data are shown as mean ±s.e.m. with n = 6 biological replicates. \*\*\**p* < 0.001, ns, not significant, calculated by unpaired t-test.

### OPTN promotes the expression of NPY to prevent RGC loss and maintain neuroimmune homeostasis in the retina

Our study demonstrated that OPTN plays a role as a neuroimmunomodulator in the pathogenesis of glaucoma. The extent to which the neuroprotective effect of OPTN extends to other retinal degenerative diseases remains uncertain. As a well-established model of retinal degeneration, the optic nerve crush (ONC) model enables us to accurately investigate the correlation between OPTN expression and optic neuropathy. Through bioinformatics of open-source databases^61^, we found that OPTN is enriched in resilient RGCs in the ONC model and that its expression levels significantly increase following optic nerve injury (Fig. 7a). This suggests that OPTN expression has the potential to protect RGCs from death. Moreover, we examined the DEGs between OPTN positive and OPTN negative RGCs in the intact group and found that the expression level of FAM19A4 and neuropeptide Y (NPY) significantly increased in the OPTN-expressed RGCs (Fig. 7b). NPY is a 36 amino acid peptide and is involved in many physiological processes, including stress response and cortical excitability. The current study shows that activation of the NPY receptor could partially rescue the RGC death in the ocular hypertension glaucoma model^62^, suggesting that OPTN may promote NPY expression to prevent RGC death from glaucoma pathology. FAM19A4 encodes the chemokine that acts as a regulator of immune and nervous cells, supporting the mediator role of OPTN in neuroimmune homeostasis in the retina. Further bioinformatic analysis on open-sourced RGC single-cell RNA sequencing (scRNA-seq) datasets^61^ reveals that NPY is significantly enriched in OPTN-positive and resilient RGCs, with increased expression after neural injury (Fig. 7c; Supplementary Fig. 5b, 7b). Moreover, based on data from POAG patients^63^, our bioinformatic analysis has demonstrated a significant downregulation of both OPTN, NPY, and FAM19A4 under glaucomatous degeneration (Fig. 7d, Supplementary Fig. 6). We further examined the expression level of NPY in retinal ganglion cell layer (GCL) and inner plexiform layer (IPL) after OPTN knockout in the retina (Fig. 7e). The expression level of NPY is significantly decreased following OPTN knockout (Fig. 7f), indicating that OPTN may achieve its neuroprotective effect by regulating NPY expression.

**Fig. 7.**
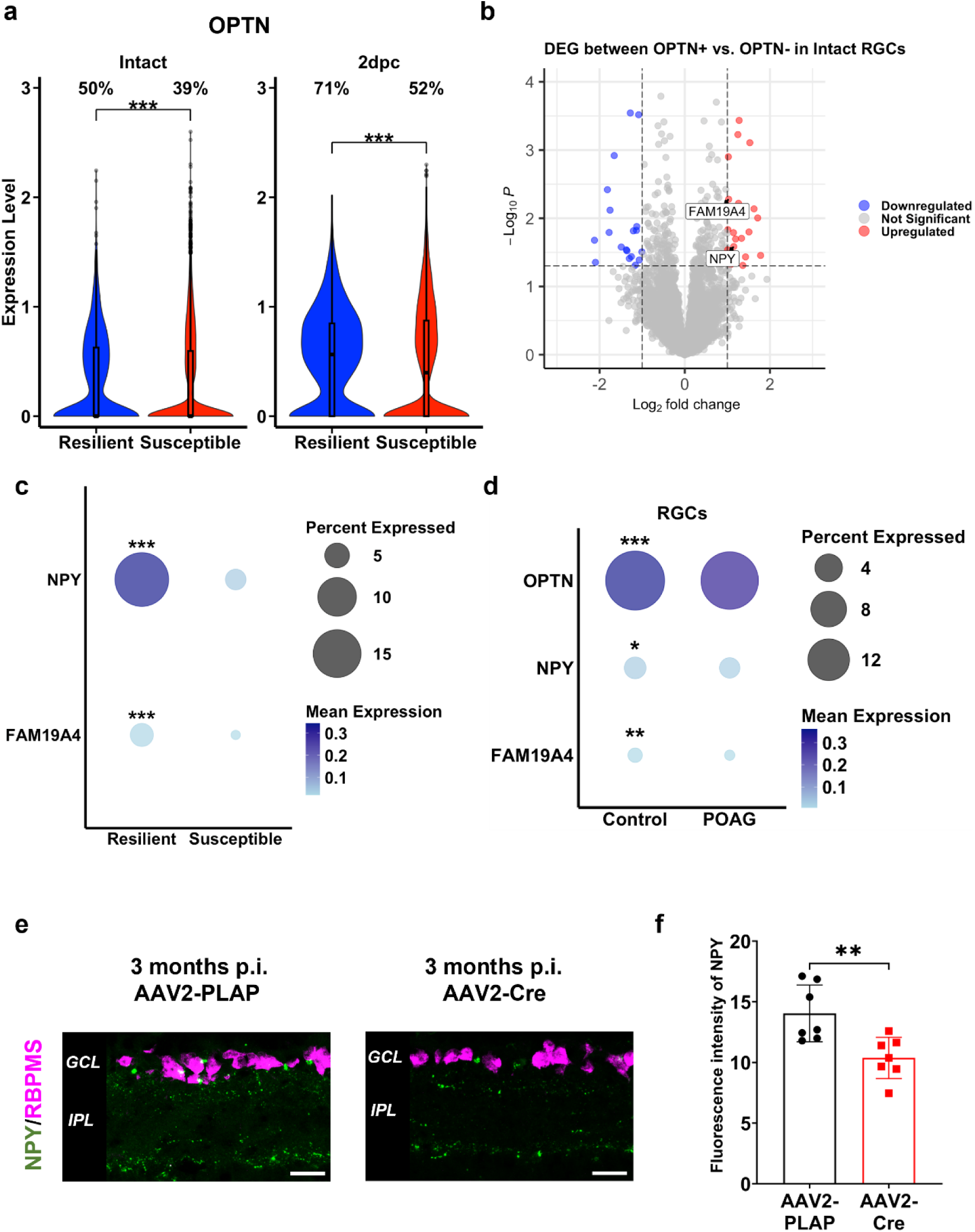
Bioinformatics analysis of scRNA-seq dataset of RGC. **a** Single cell RNA-seq analysis of OPTN in intact and injured RGCs. Res, resilient RGCs; Sus, susceptible RGCs. Numbers on top of the violin plots show the percentage of RGCs expressing the analyzed genes. **b** Volcano plot showing differentially expressed genes between OPTN+ and OPTN-intact RGCs. **c** Dot plot showing expression level and percentage expressed of NPY and FAM19A4 between Resilient and Susceptible RGCs. **d** Dot plot showing expression level and percentage expressed of OPTN, NPY, and FAM19A4 in control and POAG patients. \*\*\**p* < 0.001, \*\**p* < 0.01, \**p* < 0.05, calculated by Wilcoxon rank sum test. **e** Representative IHC images of NPY fluorescent intensity after OPTN KO. **f** Quantification of NPY fluorescent intensity after OPTN KO. \**p* < 0.05.

Previous genome-wide sequencing studies have demonstrated the activation of CHOP and ATF3 in retinal neurodegeneration, both of which target neuroinflammatory pathways^43^. Given that OPTN knockout induces neuroinflammation, we subsequently investigated whether it activates the CHOP and ATF3 expression (Supplementary Fig. 8a, 9a). Our results demonstrated that the CHOP signaling pathway is activated following OPTN loss-of-function but not ATF3 (Supplementary Fig. 8b, 9b). It implies that these injury-induced transcription factors may have different regulatory mechanisms in retinal neurodegeneration. These findings suggest that the OPTN-NPY signaling pathway may suppress the CHOP-mediated neuroinflammation to maintain neuroimmune homeostasis in the retina.

## Discussion

One major finding of our study is the critical role of wild-type OPTN in RGC survival and retinal neuroimmune homeostasis. OPTN loss of function contributes to retinal gliosis, neuroinflammation and RGC degeneration, which is complementary to the widely recognized gain-of-function mechanisms of OPTN. Gain-of-function OPTN mutations such as E50K or M98K have been demonstrated to induce RGC degeneration by enhancing the OPTN-TBK1 pathway in NTG^16,64,65^. Our results are consistent with the implicated roles of OPTN loss-of-function mutations in ALS^66^, implicating that OPTN mutations may contribute to optic neuropathies in both loss-of-function and gain-of-function manner. Interestingly, recent studies investigating the role of endogenous OPTN dysfunction in neurodegeneration have yielded divergent findings. Consistent to our results, by employing the mouse γ-synuclein (mSncg) promoter to specific knockout OPTN in RGCs, one recent study has demonstrated that OPTN dysfunction results in IOP-independent RGC death and axon degeneration by decreasing axonal mitochondrial transport^53^. Given that AAV2 may also transduce amacrine cells^67^, we cannot entirely exclude the possibility of non-cell-autonomous RGC death mechanisms induced by OPTN loss-of-function. Intriguingly, another recent study has suggested a reduced total thickness of the ganglion cells and inner plexiform layers, while the RGC loss was not statistically significant in a mouse model with CMV-driven systemic knockout of OPTN from first coding exon^68^. It should be noted that our study flanked exon 12 of OPTN by LoxP sites, resulting in a version of OPTN with C-terminus truncation, which has been found in both NTG^8^ and ALS^69^ patients. Intriguingly, OPTN N-terminal coiled-coil domain (NTD), within which gain-of-function mutations such as E50K and M98K have been identified, is responsible for the OPTN/TBK1^64,70^. It would be interesting to hypothesize that mutations at different domains of OPTN may result in different pathological mechanisms. In addition, more binding partners of OPTN/TBK1 complex might also contribute to neuroprotection or neurodegeneration, which certainly awaits more future investigations.

Neuroinflammation is a critical hallmark of many neurodegenerative conditions, such as Alzheimer’s disease^71^, Parkinson’s disease^72^, Huntington’s disease^73^, and ALS^74^. In the case of optic neuropathies such as glaucoma, activation of microglia and astrocytes is recognized as a relatively initial events of neural damage preceding RGC loss^75–77^. Our study suggests that OPTN loss-of-function may contribute to neuroinflammation that exacerbates RGC death in both HTG and NTG caused by OPTN mutations. This observation aligns with the previous studies showing that inhibiting microglia activation can delay RGC death^40,78^. Using our optimized viscobead injection model, we found that OPTN loss of function does not further exacerbate RGC degeneration in ocular hypertension. This observation suggests that OPTN deficiency might affect an overlapped subset of vulnerable RGCs with ocular hypertension. One limitation of our study is that we did not pinpoint the exact susceptibility of each RGC subtype to OPTN deficiency in normal tension and hypertension conditions. To precisely identify vulnerable RGC subtypes in OPTN deficiency, future single-cell transcriptomic analysis may be warranted. Our study has, for the first time, identified a shared OPTN-driven neuroprotective mechanism underlying retinal neurodegeneration in both NTG and HTG. Previous studies have identified CHOP and ATF3 as important injury-induced TFs that activate a set of neuroinflammation genes^43,49,50,79^. Our study has also revealed that OPTN loss of function leads to upregulation of CHOP, a major pro-apoptotic gene induced by ER stress^47,49^, which is consistent with the *in vitro* evidence that OPTN regulates the ER stress-induced signaling pathways and cell death^28^. Intriguingly, ATF3, whose expression has been demonstrated to be drastically increased in acute neural injury^43,79^, was not activated in OPTN knockout condition (Supplementary Fig. 9). One possible explanation is that the gradual loss of RGCs in our normal tension optic neuropathy model did not reach the threshold for the injury induced up-regulation of ATF3 that can be observed in ONC injury^43,61,80^.

ER stress is closely linked to microglial activation^81^ and has been reported in both NTG and HTG models^24,25^. Our bioinformatic analyses and IHC validation have revealed that NPY enriched in OPTN-expressing RGCs and OPTN knockout results in the downregulation of NPY, while NPY has been demonstrated to inhibit ER stress^82,83^. Furthermore, many studies have demonstrated that NPY functioning as an immunomodulator represents a neuroprotection in glaucomatous degeneration^62,84,85^. These findings suggest that OPTN dysfunction-induced downregulation of NPY may disinhibit ER stress and result in persistent microglia activation and progressive RGC death. Our study reveals how NPY interacts with ER stress within an OPTN-mediated pathway, which is crucial for maintaining neuroimmune homeostasis. This discovery highlights the OPTN-NPY pathway as a novel and promising therapeutic target for NTG, HTG, and potentially other neurodegenerative conditions, especially for cases irresponsive to IOP-lowering interventions.

Anti-inflammatory treatments for neurodegeneration have yielded promising outcomes in pre-clinical and clinical studies by effectively preventing CNS neuron death in optic neuropathy^86^, Alzheimer’s Disease^87^, and ALS^88^. Nevertheless, further understanding of the intrinsic mechanisms underlying neuroimmune dysregulation in neurodegeneration is essential to optimize such therapeutic strategies. We have demonstrated an endogenous neuroprotective mechanism by wild-type OPTN and its downstream NPY, potentially minimizing off-target effects associated with broad-spectrum neuroimmune modulation. Consequently, the OPTN-driven neuroprotective mechanism, independent of IOP reduction, offers a complementary strategy to existing glaucoma treatments. In conclusion, the OPTN-driven inhibitory impact of NPY on CHOP-mediated neuroinflammation represents a promising target for neuroimmune modulation in neurodegenerative diseases.

## Methods

### Animals

All experimental procedures were performed in compliance with animal protocols approved by the IACUC at Beth Israel Deaconess Medical Center and Harvard Medical School. OPTN floxed mice or wild-type control mice aged 4 weeks were used for OPTN knockout through intravitreal AAV2 injection. Viscobead injection was conducted at the age of 8 weeks. Male and female mice were used in this study at ratios dependent on litter available and with equal distributions across experiments conducted extemporaneously. OPTN floxed mouse strain (*Optn^tm1.1Jda^*/J) was obtained from Jackson Laboratory (Stock# 029708).

### Intravitreal AAV injection

For all surgical procedures, mice were anesthetized with ketamine and xylazine and received Buprenorphine as a postoperative analgesic. As previously described, intravitreal virus injection was performed at the age of 4 weeks. Briefly, a pulled-glass micropipette was inserted near the peripheral retina behind the ora serrata and deliberately angled to avoid damage to the lens. 2 μl of the AAV2/2-CAG-Cre virus was injected for OPTN fl/fl mice. For OPTN knockout injection, the titer of each AAV2-CAG-Cre was adjusted to 1 × 10^12^ genome copies/mL.

### Viscobead-induced experimental mouse glaucoma model

The elevation of IOP was induced by injection of viscobead to the anterior chamber of mouse eyes. The surgery procedures were modified by a well-established microbead occlusion model^89^. Briefly, by using a standard double emulsion method, poly-d,l-lactic-co-glycolic acid (PLGA) / polystyrene (PS) core-shell microparticles (viscobead) with 1-20 um size distributed were first fabricated at a concentration of 30% (v/v) in saline. The corneas of anesthetized mice were gently punctured near the center using a 33g needle (CAD4113, sigma). A bubble was injected through this incision site into the anterior chamber to prevent possible leakage. Then, 1 μL viscobead was injected into the anterior chamber. After 5 min when the viscobead accumulated at the iridocorneal angle, the mouse was applied antibiotic vetropolycin ointment (Dechra Veterinary Products, Overland Park, KS) and placed on a heating pad for recovery.

### Intraocular pressure measurement

The IOP measurements were performed using a TonoLab tonometer (Colonial Medical Supply, Espoo, Finland) according to product instructions. Mice were first anesthetized with a sustained isoflurane (NDC 14043-704-05, Patterson Veterinary) flow (3% isoflurane in 100% oxygen). The administration of isoflurane was discontinued once the mice were anesthetized, and IOP measurements were conducted within a 3-minute timeframe, in order to minimize the influence of isoflurane on IOP. Average IOP was generated automatically with five measurements after the elimination of the highest and lowest values.

### Perfusions and tissue processing

For immunostaining, animals were given an overdose of anesthesia and transcardiacally perfused with ice-cold PBS followed by 4% paraformaldehyde (PFA, sigma). After perfusion, retinas were dissected out and postfixed in 4% PFA overnight at 4⁰C. Tissues were cryoprotected by sinking in 30% sucrose in PBS for 48 hours. Samples were frozen in Optimal Cutting Temperature compound (Tissue Tek) using dry ice and then sectioned at 14 μm for retinas.

### Retinal wholemount staining and quantification of RGC survival

Dissected retinas were rinsed in PBS and then blocked in PBS with 1% Triton X-100 and 5% horse serum (wholemount buffer) overnight at 4 °C. Retinas were then incubated with primary antibodies diluted in wholemount buffer for 2-4 days at 4 °C, followed by three rinses with PBS (10 min each time). Next, retinas were incubated with secondary antibodies (all with 1:500 dilution) diluted in PBS overnight at 4 °C. Finally, after five PBS washes (10 min each time), retinas were mounted with Fluoromount-G (Southern Biotech, Cat. No. 0100-01).

### Immunostaining and imaging analysis

Cryosections (14-μm thick) were permeabilized and blocked in blocking buffer (0.5% Triton X-100 and 5% horse serum in PBS) for 1 hour at room temperature and overlaid with primary antibodies against OPTN, RBPMS, Iba1, CD68, and GFAP (OPTN, Cayman, No. 100000, 1:100; RBPMS, Raygene, A008712, 1:500; Iba1, Novus, NB100-1028, 1:500; CD68, Biorad, MCA1957T, 1:500; GFAP, Dako, Z0334, 1:500) overnight at 4 °C. On the next day, the corresponding Alexa Fluor 488-, 555- or 647-conjugated secondary antibodies were applied (all secondary antibodies were purchased from Invitrogen). All stained sections were mounted with solutions with DAPI-containing mounting solution and sealed with glass coverslips. All immunofluorescence-labeled images were acquired using Zeiss 700 or Zeiss 710 confocal microscope. For each biological sample, 3-5 sections of each retina were imaged and were taken under 10x or 20x objectives for quantification. Positive cell numbers were then quantified manually using the Plugins/ Analyze /Cell Counter function in ImageJ software.

### scRNA-seq analysis – differential gene expression

The *Homo sapiens* eye dataset was obtained from Human Cell Atlas and only the normalized UMI matrix was used for further analysis^63^. The RGC dataset GSE137398 underwent a standard Seurat pipeline with the R package Seurat^90^. Differential expression analysis was conducted with the Bioconductor package DESeq2^91^. Statistical significance of differentially expressed genes (DEGs) was determined at padj < 0.05 and |Log2FC| > 1. Statistical significance between groups in scRNA-seq dataset was determined using the nonparametric alternative of the t-test, the Wilcoxon rank-sum test.

### Statistical analysis

The normality and variance similarity were measured by Microsoft Excel and R programming Language before we applied for any parametric tests. A two-tailed student’s t-test was used for the single comparison between the two groups. The rest of the data were analyzed using one-way or two-way ANOVA depending on the appropriate design. *Post hoc* comparisons were carried out only when the primary measure showed statistical significance. The P-value of multiple comparisons was adjusted by using Bonferroni’s correction. Error bars in all figures represent mean ± S.E.M. The mice with different litters, body weights and sexes were randomized and assigned to different treatment groups, and no other specific randomization was used for the animal studies.

## Data Availability

The authors declare that the data supporting the findings of this study are available within the article and its Supplementary Information files, or are available upon reasonable requests to the authors.

## Acknowledgments

We sincerely thank Dr. Zhigang He from Boston Children’s Hospital for constructive suggestions. This work was supported by grants from the National Institutes of Health (EY032181).

## Authors’ contributions

Qinglong Wang, Y. W., R.D. and F.T. conceived and Qinglong Wang., Y.W., Y.D.D.J., G.C. performed the experiments and analyzed the data. Qinglong Wang., Y.W. and G.C. performed sample collection. Y.D.D.J performed bioinformatic analyses. Qinglong Wang, Y.W., Y.D.D.J., G.C., W.Y., H.G., J.H., Y.L., Qianbin Wang, and F.T. prepared the manuscript with the input from all authors.

## Additional Information

Correspondence and requests for materials should be addressed to Feng Tian.

